# Craft: A Machine Learning Approach to Dengue Subtyping

**DOI:** 10.1101/2025.02.10.637410

**Authors:** Daniel J. van Zyl, Marcel Dunaiski, Houriiyah Tegally, Cheryl Baxter, The INFORM Africa research study group, Tulio de Oliveira, Joicymara S. Xavier

## Abstract

**Motivation:** The dengue virus poses a major global health threat, with nearly 390 million infections annually. A recently proposed hierarchical dengue nomenclature system enhances spatial resolution by defining major and minor lineages within genotypes, aiding efforts to track viral evolution. While current subtyping tools – Genome Detective, GLUE, and NextClade – rely on computationally intensive sequence alignment and phylogenetic inference, machine learning presents a promising alternative for achieving accurate and rapid classification.

**Results:** We present Craft (**C**haos **Ra**ndom **F**ores**t**), a machine learning framework for dengue subtyping. We demonstrate that Craft is capable of faster classification speeds while matching or surpassing the accuracy of existing tools. Craft achieves 99.5% accuracy on a hold-out test set and processes over 140 000 sequences per minute. Notably, Craft maintains remarkably high accuracy even when classifying sequence segments as short as 700 nucleotides.

**Supplementary information:** A supplemental table acknowledging the authors of the GISAID dengue sequences is available at *Bioinformatics* online.

## Introduction

Dengue is a systemic viral infection transmitted primarily by Aedes mosquitoes that poses a significant global health threat. Primarily affecting tropical and subtropical regions, dengue can lead to severe illness and even death, with approximately 390 million infections occurring annually (Bhatt et al., 2013).

The traditional classification of the dengue virus is structured around four primary serotypes, each subdivided into distinct genotypes. Initially, genotypes were defined by a pairwise genetic distance greater than 6% within a 240 nucleotide sequence of the envelope (E) coding region (Rico-Hesse, 1990). With the accumulation of more extensive sequence data, this classification expanded to incorporate entire protein-coding regions. Many of these genotypes were established two to three decades ago (Twiddy et al., 2003; Lanciotti et al., 1994). This traditional framework has played a pivotal role in capturing the transmission dynamics of the dengue virus. However, enhanced global sequencing capacity, coupled with its integration into public health systems, has the opportunity to achieve finer granularity in classifying the diversity of dengue.

Hill et al. (2024) proposed a refined dengue virus nomenclature system that introduces additional classification levels, termed major and minor lineages, within each genotype. This addresses key limitations of the traditional genotype-based classification system, including its inability to fully capture the genetic diversity of the virus and the growing demand for finer spatial resolution.

The proposed nomenclature draws inspiration from the Pango system, a hierarchical lineage system designed to track the evolution of SARS-CoV-2 (Rambaut et al., 2020), as well as systems implemented for rabies (Campbell et al., 2022) and mpox viruses (Happi et al., 2022). Alongside their lineage designation system, Hill *et al*. Hill et al. (2024) referred the reader to three publicly available subtyping tools that could be used to classify sequences according to the new nomenclature; namely Genome Detective, GLUE, and Nextclade.

The Genome Detective subtyping tool (Fonseca et al., 2019) performs lineage classification in a two-step process: species identification and clade assignment. During species identification, the serotype is identified using BLAST (Altschul et al., 1990) to generate up to three potential matches. Advanced Genome Aligner (AGA) (Deforche, 2017) refines these matches by computing overlap and concordance scores. The sequence is assigned to the reference with the highest product of these scores. During clade assignment, maximum likelihood phylogenies are constructed using IQ-TREE 2 (Minh et al., 2020). The Genome Detective dengue typing tool is available as a free web application^1^.

GLUE (Singer et al., 2018) employs Maximum Likelihood Clade Assignment (MLCA) for genotyping, leveraging reference phylogenies for robust sequence classification. MAFFT (Katoh et al., 2002) integrates query sequences into the multiple sequence alignment. The RAxML Evolutionary Placement Algorithm (Stamatakis, 2014) determines the optimal placement of the query within a fixed reference tree. During clade assignment, the GLUE engine interprets phylogenies to classify sequences based on evolutionary distance and topological relationships relative to reference sequences. GLUE is available as a free local command-line tool^2^.

Nextclade (Aksamentov et al., 2021) assignments are based on identifying the critical substitutions that define lineages. During substitution analysis, uniquely informative nucleotide and amino acid substitutions are identified, with substitutions mapped to polyprotein-coding regions. The identification of these substitutions associated with previously classified branches is done manually to ensure accurate substitution association with the developed classification. Substitution tables are then generated following Augur Clades standards. Upon inference, these critical substitutions are then identified and used to inform classification. NextClade is available as both a free web application and a command-line tool^3^.

Genome Detective, GLUE, and NextClade, in conjunction with most current sequence classification methods, are based on sequence alignment and phylogenetic inference. Both processes are computationally expensive. Additionally, alignment-based tools often assume sequence collinearity; they expect sequences to be arranged in the same order. This assumption is frequently violated in real-world conditions, especially in viral genomes, which are characterized by high mutation rates and frequent recombination events (Zielezinski et al., 2017).

Alignment-free (AF) sequence comparison techniques, as the name suggests, do not rely on traditional sequence alignment (Zielezinski et al., 2019), instead drawing on statistical and mathematical methods to compare sequences based on their composition. Most AF techniques reduce sequences to numerical vectors representing their most informative characteristics (Bonidia et al., 2021), which can then be used to calculate pairwise distances for phylogenetic classification.

AF techniques have been more recently adopted as feature extraction techniques for machine learning applications. Cacciabue et al. (2022) developed Covidex , an AF tool leveraging *k*-mer frequency profiles as input for Random Forest classifiers, achieving 96.56% accuracy in classifying 1 437 SARS-CoV-2 Pango lineages. Building on this, INFINITy (Cacciabue and Marcone, 2023) was introduced as an AF tool for influenza virus subtyping and clade classification, also utilizing *k*-mer embeddings and Random Forest models.

Wade et al. (2024) provided a comparative analysis of sequence vectorization methods for HIV-1 subtyping and identified *k*-mer-based feature extraction combined with XGBoost as the most effective approach.

Finally, van Zyl et al. (2024) evaluated six alignment-free (AF) methods as feature extraction techniques for rapid and scalable machine learning-based viral sequence classification. They showed that word-based techniques achieve over 97% classification accuracy when distinguishing 3 502 distinct SARS-CoV-2 lineages using a dataset of 297 186 sequences. Additionally, Van Zyl *et al*. applied AF feature extraction techniques to dengue and HIV datasets and achieve near-perfect classification accuracy for dengue (serotype and genotype) and over 89% accuracy for HIV. In this paper, we introduce Craft (**C**haos **Ra**ndom **F**ores**t**)^4^, a machine learning approach for dengue subtyping under the newly proposed dengue nomenclature. Craft uses a novel feature extraction technique which modifies the Frequency Chaos Game to be bitwise in combination with a Random Forest classification model. With Craft, we are able to achieve highly competitive classification performance compared to Genome Detective, NextClade and GLUE, whilst being significantly less computationally expensive.

## Methods

The objective of this research is twofold: first, to identify a machine learning method that is computationally efficient and can achieve high classification accuracy for subtyping according to the newly established dengue nomenclature; and second, to compare both the classification accuracy and speed of this method against existing dengue subtyping tools.

### Data Curation

To train and evaluate supervised machine learning models for our experiments, we required a comprehensive dataset of complete dengue sequences along with their corresponding lineage assignments.

We sourced complete nucleotide sequences from the Global Initiative on Sharing All Influenza Data (GISAID) and the National Center for Biotechnology Information (NCBI) databases. To ensure high-quality data for reliable analysis, we only included sequences with coverage greater than 95%. No additional filters were applied, allowing the inclusion of sequences from any geographic location or collection date. This process resulted in an initial dataset comprising 19 470 complete nucleotide sequences.

For lineage assignments, we utilized Genome Detective, GLUE, and NextClade to assign lineages to the selected sequences. All three tools used the latest version of the dengue nomenclature as of December 20th, 2024. Sequences were divided into batches of 1 000, to ensure that each batch contained sequences from only a single serotype. This segmentation was necessary, as both NextClade and GLUE can only perform subtyping within a single serotype at a time. For each batch, we captured the time it took to complete the assignment process. Thus, each sequence in the dataset received three lineage assignments, one from each tool. We excluded sequences from further analysis for which any of the tools failed to provide an assignment.

The global spread of dengue has led to an uneven distribution of lineages in the available dataset, creating a risk of unintended bias in machine learning models that might subsequently favor more prevalent lineages. To mitigate the issue of class imbalance, we limited the dataset to a maximum of 100 samples per class, according to lineage assignments provided by Genome Detective. Although this approach allowed certain GLUE and NextClade classes to potentially exceed the 100-sample limit, the overlap among the three labeling tools was substantial enough for this variation to be considered negligible in the context of training machine learning models.

To further refine the dataset, we ensured that each class had a sufficient number of training samples across the assignments provided by each labeling tool. This resulted in the issue that the removal of samples due to label gaps in one tool could inadvertently impact the class distribution in another. To address this complexity, we tested various values for minimum class size threshold and ultimately selected a threshold of six samples per class per tool. This approach provided an optimal balance between ensuring sufficient training samples per class and maintaining a representative number of distinct classes. Of the initial 185 classes, only 13 were excluded for failing to meet this minimum threshold. This final refinement reduced the dataset to 8 506 sequences.

To assess the consistency of classification among the three subtyping tools in the finalized dataset, we conducted a comparative analysis of the lineage assignments provided by Genome Detective, GLUE, and NextClade. The results of this comparison are summarized in Table 1.

**Table 1.** Agreement frequency between Genome Detective, GLUE and NextClade class assignments.

| Agreement Category | Frequency |
| --- | --- |
| All Agree | 6152 times |
| Only Genome Detective & NextClade Agree | 2159 times |
| Only Genome Detective & GLUE Agree | 107 times |
| Only NextClade & GLUE Agree | 70 times |
| None Agree | 18 times |

Among the 8 506 observations, all three tools agreed on the lineage assignment in 6 152 cases. In the majority of instances where only two tools agreed on an assignment, GLUE was the outlier. Notably, there were only 18 cases where all three tools assigned completely different labels.

Given the observed disagreement among the tools, we established a consensus labeling approach for model training and evaluation. Consensus labels were determined using a majority-vote strategy: for each sample, the label assigned by at least two of the tools was selected. In the 18 cases where all three tools assigned different labels, the consensus label was derived based on the most specific evolutionary level agreed upon by at least two of the three tools. For example, if the supplied predictions were 3III B, 3III B.3, and 3III, the assigned consensus label would be 3III B, the most specific assignment for which two of the tools agree.

After establishing the consensus label set, the dataset was stratified by lineage (according to the consensus labels) and split into training and testing sets using a 5:3 ratio. Each class within the consensus set included a minimum of eight observations. This division enabled 5-fold cross-validation for model validation while preserving the integrity of the dataset for testing.

### Feature Extraction: The Bitwise Chaos Game Representation

Building on the findings of van Zyl et al. (2024), we used the Frequency Chaos Game Representation (FCGR) to extract descriptive feature vectors from the dengue sequence dataset. van Zyl et al. (2024) demonstrated that Random Forest models trained on FCGR feature vectors for both SARS-CoV-2 and dengue (serotype and genotype classifications) achieved the highest accuracy and Macro F1 scores among the feature extraction methods considered in their study.

The FCGR process begins by generating an image with resolution *r* x *r*, where *r* is a hyperparameter that determines the granularity of the representation. Each corner of the image is assigned one of the four nucleotide bases: A, C, T, or G. The input sequence is then processed character by character, starting at the center of the image. For each nucleotide read, the process moves halfway toward the corresponding corner and increments the pixel value at the resulting location by one, following the principles of the chaos game. This continues for the entire sequence, skipping ambiguous bases. The resulting image, which captures the nucleotide frequency distribution in spatial form, is then flattened into a feature vector. The FCGR vector can be computed efficiently in linear time.

In this research, we explored a modification of the FCGR. Instead of tallying pixel counts, this approach uses binary values to indicate only the presence or absence of specific pixels. This modified representation, referred to as the Bitwise Chaos Game Representation (BCGR), achieves competitive accuracy when used with Random Forest models while offering distinct advantages. BCGR simplifies the feature set, reduces model complexity, and minimizes storage requirements. The similarity between BCGR and FCGR depends on the resolution (*r*) and the length of the sequences that are analyzed. At higher resolutions (*r*), the probability of pixels being incremented multiple times in the FCGR decreases. Conversely, with longer sequences, the likelihood of pixels being incremented multiple times increases.

### Cross-Validation

We used 5-fold cross-validation on the training set to optimize the parameters for both the feature extraction techniques and the Random Forest models. For the Random Forest classifiers, we evaluated two splitting criteria: Gini impurity and entropy. Additionally, we tested two class-weighting schemes: no weighting and a balanced weighting scheme designed to mitigate class imbalance. All Random Forest models consisted of 100 trees, ensuring sufficient saturation with negligible gains from further increases.

For feature extraction, we utilized both FCGR and BCGR representations, evaluating resolution values (*r*) ranging from 32 to 256 in doubling increments. Although Random Forest models inherently perform feature selection, we improved inference efficiency by identifying the 5 000 most informative features from each trained model. We then subsequently trained Random Forests on these reduced feature sets and included them in our model validation experiments.

We conducted an extensive grid search to identify the optimal parameters for all possible configurations. The cross-validation process was performed using training labels provided by Genome Detective, GLUE, and NextClade individually, as well as the labels of the consensus set. In cases where a class had insufficient samples to support 5-fold cross-validation, we excluded that class from the training set for the corresponding experiment. This decision had minimal impact on the class distributions, affecting only five classes in the GLUE label set.

Our findings indicate that for all models, a resolution of 128, combined with balanced class weighting and the entropy splitting criterion, yields the highest validation accuracy. Table 2 summarizes the best validation accuracy scores achieved for each of the four considered Random Forest models and shows the average accuracy and standard deviation of the 5 folds.

**Table 2.** Cross-validation classification accuracy results for FCGR and BCGR when trained and validated on the different label sets. The top 5 000 features refer to the 5 000 most informative features as determined by a pre-trained Random Forest.

| Model | Feature Set | Genome Detective | GLUE | NextClade | Consensus |
| --- | --- | --- | --- | --- | --- |
| FCGR | Full | 0.989±0.003 | 0.964±0.005 | 0.995±0.001 | 0.992±0.001 |
| FCGR | Top 5000 | 0.988±0.004 | 0.964±0.006 | 0.996±0.001 | 0.992±0.003 |
| BCGR | Full | 0.987±0.005 | 0.964±0.005 | 0.995±0.001 | 0.992±0.001 |
| BCGR | Top 5000 | 0.989±0.004 | 0.963±0.006 | 0.995±0.001 | 0.993±0.003 |

Notably, all four feature extraction schemes yield very similar classification accuracy. Interestingly, the Random Forest models trained on the NextClade labels achieves the highest validation accuracy, suggesting that the labeling logic employed by NextClade is the easiest to replicate. Evaluation of the feature extraction schemes on the consensus label set reveals that the streamlined BCGR approach, which uses only the 5 000 most informative features, achieves comparable accuracy while significantly improving computational efficiency. Based on these results, we selected this model structure or the subsequent experiments on the hold-out test set. We named the chosen architecture Craft (**C**haos **Ra**ndom **F**ores**t**), alluding to the combination of a chaos-based feature representations with a Random Forest model.

Before conducting the final evaluations on the hold-out testing set, we further assessed Craft’s performance by comparing its validation results when trained on labels from each tool and then validated on each of the other tools’ labels. The findings are summarized in Table 3.

**Table 3.** Validation performance of all models on the different label sets (indicated by the four right-most column headings). The training set used for each Craft model is indicated in the “Training Labels” column.

| Model | Training Labels | Genome Detective | GLUE | NextClade | Consensus |
| --- | --- | --- | --- | --- | --- |
| Genome Detective | - | 1.000 | 0.737 | 0.977 | 0.991 |
| GLUE | - | 0.737 | 1.000 | 0.732 | 0.745 |
| NextClade | - | 0.977 | 0.732 | 1.000 | 0.986 |
| Craft | Genome Detective | 0.989±0.004 | 0.736±0.009 | 0.978±0.005 | 0.987±0.003 |
| Craft | GLUE | 0.746±0.008 | 0.963±0.006 | 0.743±0.009 | 0.754±0.010 |
| Craft | NextClade | 0.974±0.004 | 0.732±0.003 | 0.995±0.001 | 0.982±0.003 |
| Craft | Consensus | 0.984±0.004 | 0.742±0.006 | 0.983±0.003 | 0.993±0.003 |

In Table 3, we also report the accuracy scores of Genome Detective, GLUE, and NextClade when compared against each other’s labels as well as the consensus set.

Craft models trained on labels from a specific tool achieved performances closely aligned with that tool’s labeling patterns. Additionally, the Craft model trained on the consensus label set consistently achieved the second-highest accuracy when compared to the labels of each of the three subtyping tools, with the highest accuracy naturally belonging to the respective tool itself. This reinforces our choice of the consensus label set and indicates that it provides a balanced representation of the three tools.

All subsequent evaluations are based on the hold-out test set of the consensus label set.

### Classification of Sequence Segments

Accurately obtaining high coverage complete dengue sequences is often challenging due to the rapid decrease in viral load following the short viremic phase that typically precedes sample collection (Guzman et al., 2010). Although the global capacity for whole genome sequencing has improved significantly, largely due to the genomic infrastructure developed during the SARS-CoV-2 pandemic, many laboratories continue to sequence only the E coding region. This region is often prioritized as it suffices for genotyping and is faster than whole genome sequencing.

In scenarios of limited genome coverage, Hill et al. (2024) demonstrated that even partial sequences can achieve high classification accuracy. Building on this insight, we recognized the potential challenges posed by incomplete sequencing and the possibility that only specific genomic regions may be available. To address this, we extended the evaluation of classification performance beyond the commonly studied E coding region, exploring the efficacy of subtyping models when applied to various partial genomic segments. Specifically, we investigated the models’ abilities to classify sequences based on random stretches of only 700 nucleotides.

To achieve this, we trained multiple Craft classifiers, each tailored to classify sequences from a specific 500-nucleotide segment of the genome. These segments were derived from aligned sequences using NextClade, with consecutive segments overlapping by 400 nucleotides. This overlap ensured that every new segment began 100 nucleotides after the previous one, providing comprehensive coverage of the genome. Separate sets of Craft classifiers were trained for each dengue serotype, and each classifier utilized the full set of features generated by the BCGR representation, rather than the reduced set of 5 000 most informative features.

For each serotype, the corresponding NextClade reference sequence was divided into similar overlapping 500-nucleotide segments, and the BCGR representation was computed for each segment. During inference, a given 700-nucleotide sequence was analyzed by extracting its central 500 nucleotides and comparing its BCGR representation to the BCGRs of all reference segments. Using a 1-nearest-neighbor approach, the segment with the closest Hamming distance was identified, enabling accurate prediction of both the general segment location and the corresponding serotype. To improve the alignment of the inference segment with its matched reference segment, we refined the match by computing the Hamming distance between the BCGR of every possible 500-nucleotide stretch within the 700-nucleotide inference sequence and the BCGR of the matched reference segment. The alignment with the minimum Hamming distance was deemed the most optimal.

Once the inference sequence was aligned with the reference segment, the predicted match position and serotype were used to load the appropriate Random Forest model for that segment. This model was then applied to classify the sequence.

As with previous experiments, we employed 5-fold cross-validation and a grid search to identify the optimal parameter configurations for the models. For this experiment, we chose not to use the reduced feature sets, instead utilizing the full BCGR feature representation. Entropy-based splitting, balanced class weighting, and a resolution of *r* continued to deliver the best performance on the validation set.

The validation dataset was constructed by randomly selecting five 700-nucleotide segment from each sequence. To ensure sufficient sequence coverage, the starting position of these segments was restricted to a minimum of the 200th nucleotide and a maximum of the 9 500th nucleotide, as many sequences lacked sufficient data at the extreme ends. The hold-out test set was augmented in a similar manner to for the final evaluation.

## Results & Discussion

The results presented in this section are solely with regards to experiments conducted on the hold-out test set using 10 independent trials for each experiment. In each trial, 80% of the training set was randomly sampled using stratified sampling based on the consensus labels to train the model, followed by evaluation on the test set. To ensure variability in the sampled training data, different random seeds were used for each trial. This methodology allowed us to assess the model’s performance variation across different training data subsets.

Model performance was evaluated using two metrics: overall classification accuracy and Macro F1 score. While accuracy measures the proportion of correct predictions across all classes, it can be misleading in the presence of class imbalances. The Macro F1 score, which averages the F1 scores of individual classes, provides a more balanced assessment, capturing the model’s effectiveness across both majority and minority classes.

Machine learning-based approaches face a significant limitation in their reliance on extensive training datasets to achieve suitable classification performance. This poses challenges when new lineages are introduced into the nomenclature with limited available training examples. To evaluate the robustness of Craft for dengue classification, we also present performance results from a scenario where the Craft model is trained under limited data conditions, using only four samples per class per trial.

### Hierarchical Performance

Table 4 contains the classification performance results for different hierarchical levels of dengue virus classification on the hold-out testing set for each model on the consensus labels.

**Table 4.** Accuracy and Macro F1 Scores for serotype, genotype, major lineage, and minor lineage for Craft, Genome Detective, NextClade, and GLUE. Craft results are reported according average and standard deviation across 10 trials.

|  | Model | Serotype | Genotype | Major Lineage | Minor Lineage |
| --- | --- | --- | --- | --- | --- |
| Accuracy | Genome Detective | 1.0 | 0.997 | 0.992 | 0.991 |
|  | GLUE | 1.0 | 0.998 | 0.932 | 0.744 |
|  | NextClade | 1.0 | 0.999 | 0.994 | 0.986 |
| | Craft (Limited training data) | 1.0 | 0.999 | $0.986 \pm 0.003$ | $0.969 \pm 0.003$ |
|  | Craft | 1.0 | **1.000** | **$0.996 \pm 0.001$** | **$0.995 \pm 0.001$** |
| Macro F1 | Genome Detective | 1.0 | 0.980 | 0.986 | 0.989 |
|  | GLUE | 1.0 | 0.950 | 0.860 | 0.665 |
|  | NextClade | 1.0 | 0.994 | **0.991** | 0.979 |
| | Craft (Limited training data) | 1.0 | $0.994 \pm 0.001$ | $0.963 \pm 0.006$ | $0.958 \pm 0.005$ |
| | Craft | 1.0 | **$0.997 \pm 0.002$** | $0.989 \pm 0.003$ | **$0.991 \pm 0.001$** |

All four models achieve perfect accuracy at the serotypic level. Across the four classification levels, Craft consistently demonstrates the highest classification accuracy. Notably, at the genotypic and major lineage levels, NextClade achieves the second-highest accuracy. However, as classification granularity increases to the minor lineage level, NextClade’s performance (0.986) declines below that of Genome Detective (0.991). Regarding the Macro F1 score, Craft outperforms all models except at the major lineage level, where NextClade achieves a marginally higher score. Under the limited training data scenario, Craft exhibits a notable decrease in classification performance compared to the full training context. Nevertheless, it maintains high accuracy and Macro F1 scores, remaining competitive with both NextClade and Genome Detective up to the minor lineage level.

Similarly, GLUE demonstrates competitive classification performance up to the minor lineage level. However, at this level of granularity, its performance declines significantly compared to the other models.

It should be noted, that these results should be interpreted with caution and are not intended to imply that any one model is inherently more accurate than another in a general context. In the absence of an expertly constructed ground truth label set, true model accuracy cannot be definitively determined. Our findings are specific to the consensus label set used in this study. While these results primarily reflect the tendencies and agreement patterns among the models, they clearly demonstrate that Craft achieves performance highly competitive with that of Genome Detective and NextClade.

### Class-wise Performance

Figure 1 offers a detailed visual comparison of the performance of Craft, Genome Detective, and NextClade across the various dengue lineages. GLUE’s results were excluded from this analysis, given its significantly lower performance compared to the other models.

**Fig. 1:**
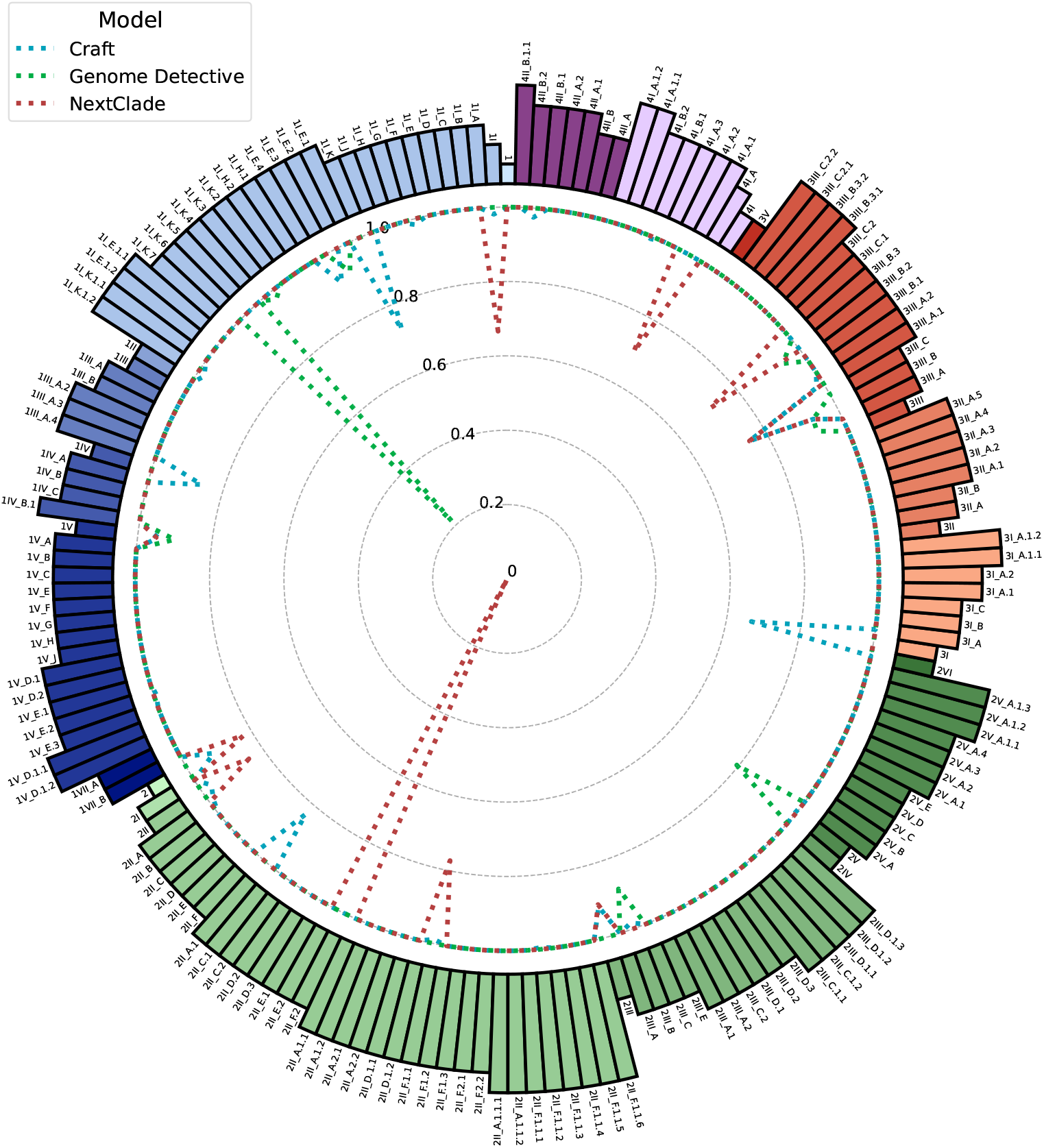
Composite radar-chart of class-wise model performance. Classes are colored according to serotype and genotype. The blue, red and green dotted lines represent Craft, NextClade and Genome Detective respectively, with lines closer to the perimeter indicating better performance. The length of each bar corresponds to the evolutionary depth of the lineage.

The class-wise performance results provide valuable insights into the behavior of each model. Notably, all three models achieve perfect performance in the majority of classes. However, for classes where this is not the case, we can observe significant performance drops, particularly for Genome Detective and NextClade. Consistent with the Macro F1 score findings, Craft demonstrates the most consistent performance across all classes.

The performance of the models shows no clear trends related to specific serotypes or hierarchical classification levels. Interestingly, the models exhibit largely distinct patterns of misclassification, rarely making the same errors across classes.

### Segment Classification

Table 5 contains the accuracy and Macro F1 scores when classifying dengue sequences based only on a random stretch of 700 nucleotides. We report the mean results achieved by Craft and their standard deviation across 10 trials.

**Table 5.** Accuracy and Macro F1 Scores for serotype, genotype, major lineage, and minor lineage across various models when classifying dengue sequences based only on a random segment of 700 nucleotides.

|  | Model | Serotype | Genotype | Major Lineage | Minor Lineage |
| --- | --- | --- | --- | --- | --- |
| Accuracy | Genome Detective | 1.000 | 0.894 | 0.782 | 0.697 |
|  | GLUE | 1.000 | 0.994 | 0.923 | 0.726 |
|  | NextClade | 0.995 | 0.974 | 0.956 | 0.921 |
| | Craft (Limited Training) | 1.000 | **$0.998 \pm 0.001$** | $0.965 \pm 0.004$ | $0.913 \pm 0.001$ |
|  | Craft | 1.000 | **$0.998 \pm 0.002$** | **$0.990 \pm 0.003$** | **$0.972 \pm 0.001$** |
| Macro F1 | Genome Detective | 1.000 | 0.923 | 0.829 | 0.748 |
|  | GLUE | 1.000 | 0.997 | 0.887 | 0.457 |
|  | NextClade | 0.998 | 0.949 | 0.972 | 0.930 |
| | Craft (Limited Training) | 1.000 | $0.989 \pm 0.090$ | $0.942 \pm 0.040$ | $0.932 \pm 0.080$ |
|  | Craft | 1.000 | **$0.994 \pm 0.040$** | **$0.991 \pm 0.030$** | **$0.972 \pm 0.040$** |

Across all metrics and classification levels, Craft consistently outperforms the other models. All models exhibit a marked decline in classification performance compared to full-genome subtyping, with this decrease being most pronounced for Genome Detective, which achieves an accuracy of only 0.696 at the minor lineage level. At this most granular level, NextClade demonstrates the second-highest performance among the models. However, NextClade also shows instances of errors at the serotypic level, indicating notable deviations from the consensus labels.

Unlike the complete sequence performance results, these findings can be interpreted more strongly, as the performance degradation of each model is clear, even without ground truth labels. With this in mind, Craft’s performance shows only minor degradation. Furthermore, when applied to a limited training scenario, Craft still achieves over 90% classification accuracy at the minor lineage level, very closely competing with NextClade, while achieving a higher Macro F1 score at that level.

Figure 2 illustrates the average classification accuracy of Craft, Genome Detective, GLUE, and NextClade across different regions of the dengue genome, measured by the starting positions of 500 nucleotide segments.

**Fig. 2:**
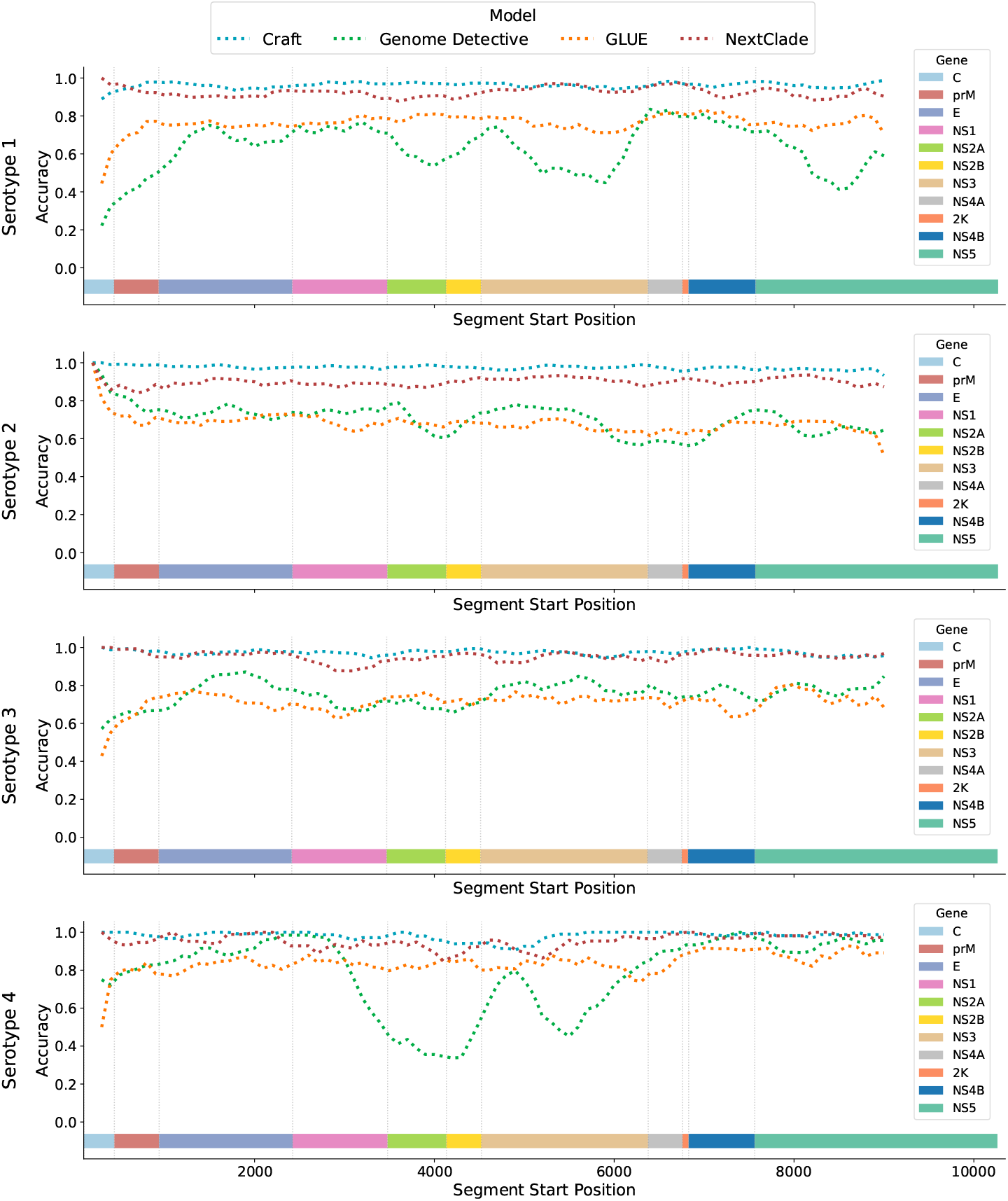
A collection of line plots showing the accuracy of each model when tested on short genomic segments from various positions within the dengue genome. Each plot corresponds to a particular serotype. We include horizontal bars indicating the positions of each gene region for each serotype.

Across most genomic regions, Craft demonstrates the highest average classification accuracy. Additionally, both Craft and NextClade exhibit lower performance variation across different regions than GLUE and Genome Detective. Variability in model performance indicates that certain genomic regions are more informative for lineage assignment, though instances of simultaneous performance drops across all models are rare. Notably, in the region surrounding the ‘E’ gene, which is the most commonly sampled segment in lieu of the whole genome, Craft and NextClade consistently achieve higher accuracy than GLUE and Genome Detective across all four serotypes.

### Classification Throughput

An additional advantage of AF feature extraction, particularly BCGR combined with Random Forest classification is its superior classification throughput and resource efficiency compared to traditional alignment-based methods and phylogenetic inference. To demonstrate this, we compared the classification throughput of Craft with Genome Detective, NextClade, and GLUE.

For Genome Detective, throughput measurements were obtained using its online web application. Craft and GLUE were benchmarked locally on a system with 24 GB of RAM and an Apple M4 Pro processor. For NextClade, throughput was evaluated both via its online web application and a local computation setup. We calculated the average and standard deviation of classification throughput for each model using the complete set of 19 470 dengue sequences. These sequences were processed in batches of 1 000, with each batch containing sequences exclusively from a single serotype. Table 6 lists the relevant results.

**Table 6.** The average throughput of Genome Detective, GLUE, NextClade and Craft in sequences per minute from 19 independent runs.

| Model | Classification Throughput |
| --- | --- |
| Genome Detective | 4.628 ± 1.505 seq/min |
| GLUE | 63.25 ± 3.51 seq/min |
| Nextclade (Local) | 7 618.98 ± 1 872.17 seq/min |
| Nextclade (Online) | 7 067.87 ± 1 430.61 seq/min |
| Craft | 141 183.70 ± 1 599.35 seq/min |

The recorded timings should not be considered exact due to differences in computational setups and conditions, however, they provide a reasonable approximation of classification throughput of the various methods. Among the models, Genome Detective is by far the slowest, with an average throughput of just 4.628 sequences per minute. In contrast, NextClade, despite being an alignment-based method, achieves a significantly higher throughput, averaging 7 067.87 sequences per minute. The efficiency advantage of alignment-free subtyping tools becomes particularly evident with Craft achieving an extraordinary throughput of over 140 000 sequences per minute.

While the current volume of dengue sequences may not fully exploit the speed advantage of Craft over NextClade, its potential becomes clear when considering the rapid growth of next-generation sequencing capabilities. The potential benefits of near real-time tracking become particularly significant in epidemic scenarios, such as with the SARS-CoV-2 pandemic, where millions of genomes were sequenced and analyzed.

## Conclusion

In this paper, we introduced Craft (**C**haos **Ra**ndom **F**ores**t**), which is a novel alignment-free machine learning framework for dengue virus subtyping under the newly proposed hierarchical nomenclature system. This new nomenclature introduced major and minor lineage classifications within genotypes, which enables more granular tracking of dengue virus evolution and spread.

By leveraging a modified Bitwise Chaos Game Representation (BCGR) for feature extraction, Craft achieves competitive classification accuracy while significantly outperforming alignment-based tools such as Genome Detective, GLUE, and NextClade in terms of computational efficiency. Craft achieves 99.5% accuracy on a majority-vote consensus label set, and is able to perform to a high standard when trained on only four examples per class. Craft particularly excels in scenarios involving only short genome segments, in which it markedly outperforms current tools.

## Supporting information

GISAID Supplemental Table

## Acknowledgments

This work was supported by grants from the INFORM Africa project through IHVN (U54 TW012041), and the eLwazi Open Data Science Platform and Coordinating Center (U2CEB032224). This work was also financed in part by the Coordenação de Aperfeiçoamento de Pessoal de Nível Superior – Brasil (CAPES) – Finance Code 001.

We gratefully acknowledge all data contributors, i.e., the Authors and their Originating laboratories responsible for obtaining the specimens, and their Submitting laboratories for generating the genetic sequence and metadata and sharing via the GISAID Initiative, on which this research is based.

## Footnotes

1 https://www.genomedetective.com/app/typingtool/dengue/

2 https://github.com/giffordlabcvr/Dengue-GLUE

3 https://clades.nextstrain.org

4 Craft is being used by the Fast tool to classify Dengue. https://fast.pathotrack.health

