## Supplementary material for "Craft: A Machine Learning Approach to Dengue Subtyping": GISAID Supplemental Table

### Data Availability

GISAID Identifier: EPI_SET_250123rb
doi: 10.55876/gis8.250123rb

All genome sequences and associated metadata in this dataset are published in GISAID's EpiArbo database. To view the contributors of each individual sequence with details such as accession number, Virus name, Collection date, Originating Lab and Submitting Lab and the list of Authors, visit 10.55876/gis8.250123rb

### Data Snapshot

- EPI_SET_250123rb is composed of 6,460 individual genome sequences.
- The collection dates range from 1944-01-01 to 2024-04-29;
- Data were collected in 94 countries and territories;
- All sequences in this dataset are compared relative to the official reference sequence employed by GISAID.
